# PlasmidHunter: Accurate and fast prediction of plasmid sequences using gene content profile and machine learning

**DOI:** 10.1101/2023.02.01.526640

**Authors:** Renmao Tian, Behzad Imanian

## Abstract

Plasmids are extrachromosomal DNA found in microorganisms. They often carry beneficial genes that help bacteria adapt to harsh conditions, but they can also carry genes that make bacteria harmful to humans. Plasmids are also important tools in genetic engineering, gene therapy, and drug production. However, it can be difficult to identify plasmid sequences from chromosomal sequences in genomic and metagenomic data. Here, we have developed a new tool called PlasmidHunter, which uses machine learning to predict plasmid sequences based on gene content profile. PlasmidHunter achieved high accuracies (up to 96.7%) and fast speeds in benchmark tests, outperforming other existing tools.

## Background

Plasmids are extrachromosomal and transmissible segments of naked, double-stranded DNA that, unlike viruses, replicate autonomously within a host cell. They are common in eubacteria, but they are also found in archaea and eukaryota. Plasmids are typically circular and often much smaller than chromosomes, but their sizes vary considerably (from about 1 Kbp to over 1 Mbp) [1–3].

As agents of horizontal gene transfer (HGT) between bacterial species [4], plasmids spread the traits that might influence the characteristics, survival and fitness of the individual host, the species and the microbial communities, and thus they play an important role in the bacterial evolution and ecology. Plasmids carry non-essential and sometimes beneficial genes that help their hosts tolerate and survive hostile conditions in a changing environment. For example, a plasmid-borne gene encodes the mercuric reductase that converts toxic Hg^2+^ to a volatile and less toxic metallic Hg^0^ [5]. We now know that plasmids play a key role in spreading the antimicrobial resistance (AMR) amongst the related bacterial species [6–8], such as those conferring resistance to many commonly used antibiotics such as tetracycline and penicillin or to β-lactams (*bla*) [9] and aminoglycosides (*aad* and *aac*) [10]. In addition to transferring AMR to other bacteria, plasmids can transmit and spread other traits [4] such as virulence, toxicity and pathogenicity to a wider group of bacteria, and consequently they pose a threat to animal and human health in more than one way. The AMR, multidrug resistance (MDR), and the rise of ‘superbugs’ are all related and considered as a dire global threat to public health [11,12]. At the same time and paradoxically, they have become a valuable tool in molecular cloning, genetic engineering, gene therapy and drug production, all of which are being explored in order to improve our health and environment [13]. Plasmid-borne genes, especially the gene clusters for secondary metabolites, are now routinely used for the development of natural products and new drugs [14] including new antibiotics [15]. In addition, plasmids harbor genes that can be utilized in environmental conservation, for example, in bioremediation [16]. Given the ubiquity of plasmid-harboring bacteria in the environment including food, water, soil and even air, investigating plasmids is increasingly important to human health and environmental conservation.

Studying plasmids have recently benefited greatly from the high-throughput sequencing (HTS) of the bacterial whole genomes and microbial metagenomics. By using HTS, we have learned a great deal more about plasmids, the extent and nature of their threats to human health and their many practical uses. However, using HTS in plasmid studies is not free of challenges. For example, it is difficult to discern plasmid sequences from those of the chromosomes in the big datasets that are produced by all the existing HTS sequencers. Even after assembling the raw reads in larger contigs, the challenge remains because these assemblies usually contain many plasmid-sized contigs that have a chromosomal origin. It is, thus, crucial to develop reliable tools to distinguish plasmid sequences from chromosomal sequences in the pool of millions of reads that are resulted from sequencing of an axenic or an environmental sample.

Several methods have exploited the discernable sequence features and gene contents in plasmids versus chromosomes to develop such tools. In recent years, machine learning (ML) has been added to the list of these methods, resulting in the improvement of the new plasmid identification tools. ML algorithms take a subset of the data called the ‘training data’ as an input and examines and evaluates the correlation between ‘feature variables’ and ‘target variables,’ a process called ‘learning,’ in order to predict the ‘target variables’ based on the features of the new data. These tools differ in the features they exploit and/or the algorithm they use for modeling, and some perform better than others. For example, PlasFlow [17] uses the sequence signatures of k-mer (3 – 7 nt) frequency in the assembled contigs as the main feature to predict plasmids. With a test dataset of contigs of 1 – 1570 Kbp, it achieves an accuracy of 89.5%. Deeplasmid [18] uses the features from both sequence signatures and gene content, including GC content, homopolymer, plasmid replication origin, coding density, contig length, hit to plasmid proteins and hit to a curated Pfam database. Using a test dataset with contigs of 1 – 330 Kbp, Deeplasmid achieves an accuracy of 84.2% and an area under the receiver operating characteristic (ROC) curve (AUC) of 89.9%. The Deeplasmid’s precision is high (up to 94.5%) but its recall is relatively low (75.6%). This means that a high percentage of the true plasmids (24.4%) are not detected at all, and from all the predicted plasmids, many (5.5%) are predicted incorrectly. Of the remaining tools, PlasClass [19] uses logistic regression classifiers and a k-mer-based (3 – 7 nt) feature vector for fragments; PlasmidVerify [20] employs Naïve Bayesian classifier and the gene content of cyclocontigs; and PlasForest [21] uses a homology-based random forest and a few other sequence features. These tools have reported higher or lower accuracies and speeds using their own test datasets.

Despite the availability of these tools for plasmid detection, several issues were needed to be addressed. Firstly, the overall accuracies, recalls and precisions of these tools were not satisfactory enough to identify plasmid sequences in the assembly files with high confidence. Secondly, the accuracies of the previous tools were not made comparable using the same test dataset. Thirdly, the running time required for some of these tools were too long (up to hours).

Here, we present a new plasmid identification tool, PlasmidHunter, that uses gene content profile alone as the feature to predict plasmid sequences with no reliance on the raw sequence data, sequence topology and coverage or assembly graph. Thus, the input data for PlasmidHunter is simply any assembled sequence file produced by any modern high-throughput sequencer and assembled by any algorithm. Using the same dataset, we also demonstrate that PlasmidHunter achieves both higher accuracy (96.7%) and recall (95.1%) with reasonable speed (< 8 minutes) in comparison to the previous tools. We also present a benchmarking of all the top tools using the same test datasets of contigs with different lengths.

## Results

### Database construction

In order to build a database for gene content profiling, 36 million unique protein sequences of 25,898 complete prokaryotic genomes from NCBI RefSeq database were downloaded and processed. After clustering and singleton removal, 0.96 million representative proteins were acquired and used to populate a database for gene content profiling used in the subsequent modeling and prediction steps (**Figure 1A**).

**Figure 1.**
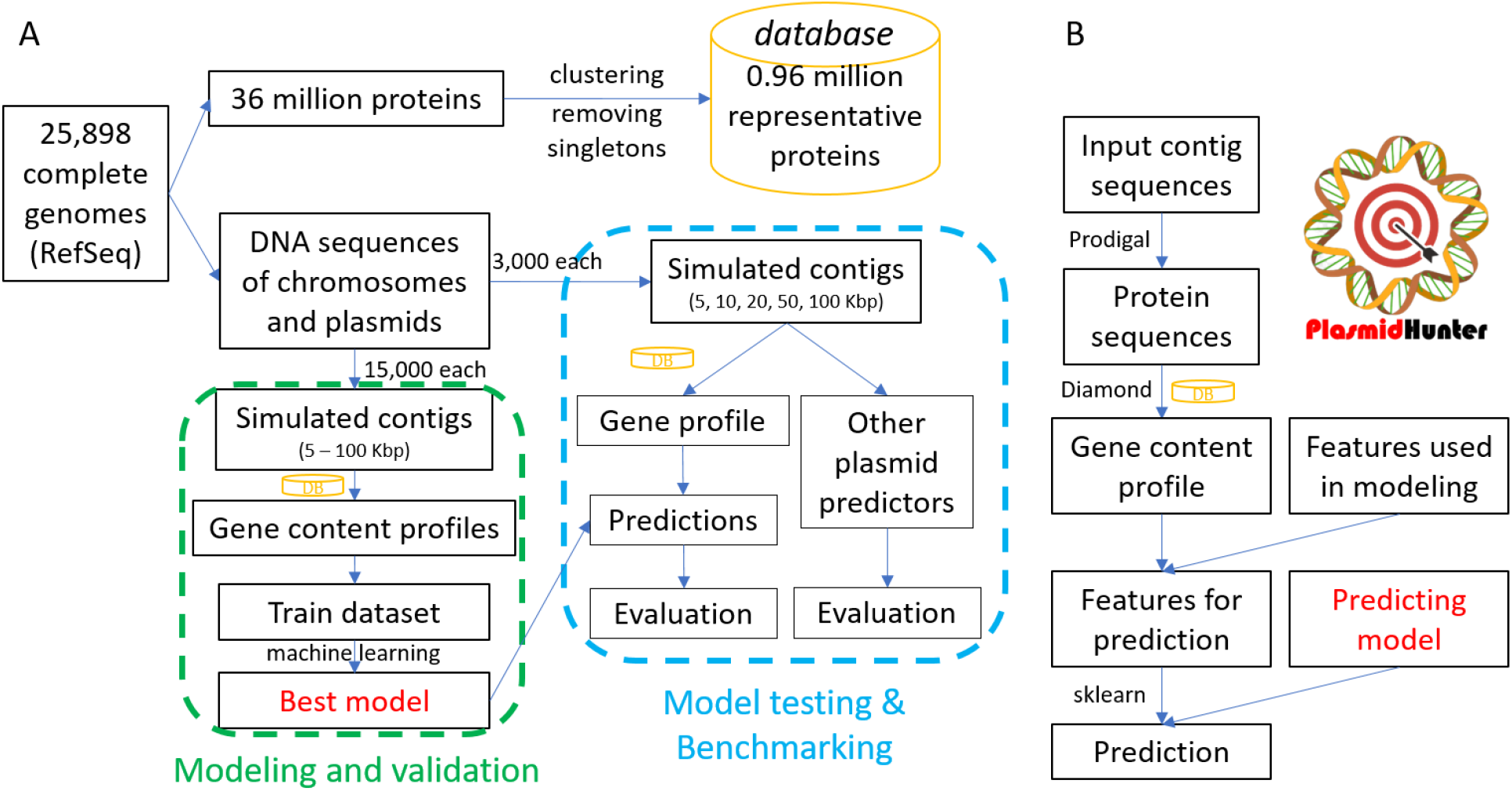
The workflow of the modeling, benchmarking and pipeline construction. (**A**) The workflow of the database construction, modeling and validation, and the model testing and benchmarking of this study. Briefly, protein sequences of 25,898 complete genomes from NCBI RefSeq database were used and 0.96 million representative protein sequences were indexed as a database. From the complete genomes, 15,000 chromosome and 15,000 plasmid sequences were used for modeling and validation, respectively, and the remaining sequences were held back for an unbiased model testing and benchmarking. The yellow cylinders represent the database. (**B**) The workflow of the pipeline PlasmidHunter. Input DNA sequences are first used to predict coding sequences of genes for each contig. The translated protein sequences are then used for Diamond alignment using the customized database. The gene content profile is filtered to retain only the gene features used in the modeling. The predicting model is then used to predict the chromosomal or plasmid origin of the contigs with the Python package sklearn.

### Gene content profile was a better discriminator of chromosome and plasmid sequences than k-mer frequency profile

In order to compare the discriminative power of gene content profile and k-mer frequency profile as the features of our model, we performed principal component analysis (PCA) on three type species, *Escherichia coli*, *Bacillus subtilis* and *Pseudomonas aeruginosa*, all with a complete genome and a plasmid (**Figure 2**). The PCA results for the k-mer frequency profile (4 – 6 bp) showed that the contigs originating from plasmids and chromosomes of different species were scattered with occasional intersections or overlaps in different regions, making It difficult to visually distinguish plasmid contigs from chromosomal contigs (**Figure 2A, B & C**). In contrast, in the PCA results for the gene content profile, all the chromosomal contigs were well separated from all the plasmid contigs. The chromosomal contigs of all the three species were highly concentrated in a single region (**Figure 2D & E**). These results clearly demonstrated the higher discriminative power of gene content profile in discerning plasmid and chromosomal sequences than that of the k-mer frequency profile. Thus, we chose gene content as the feature for our modeling.

**Figure 2.**
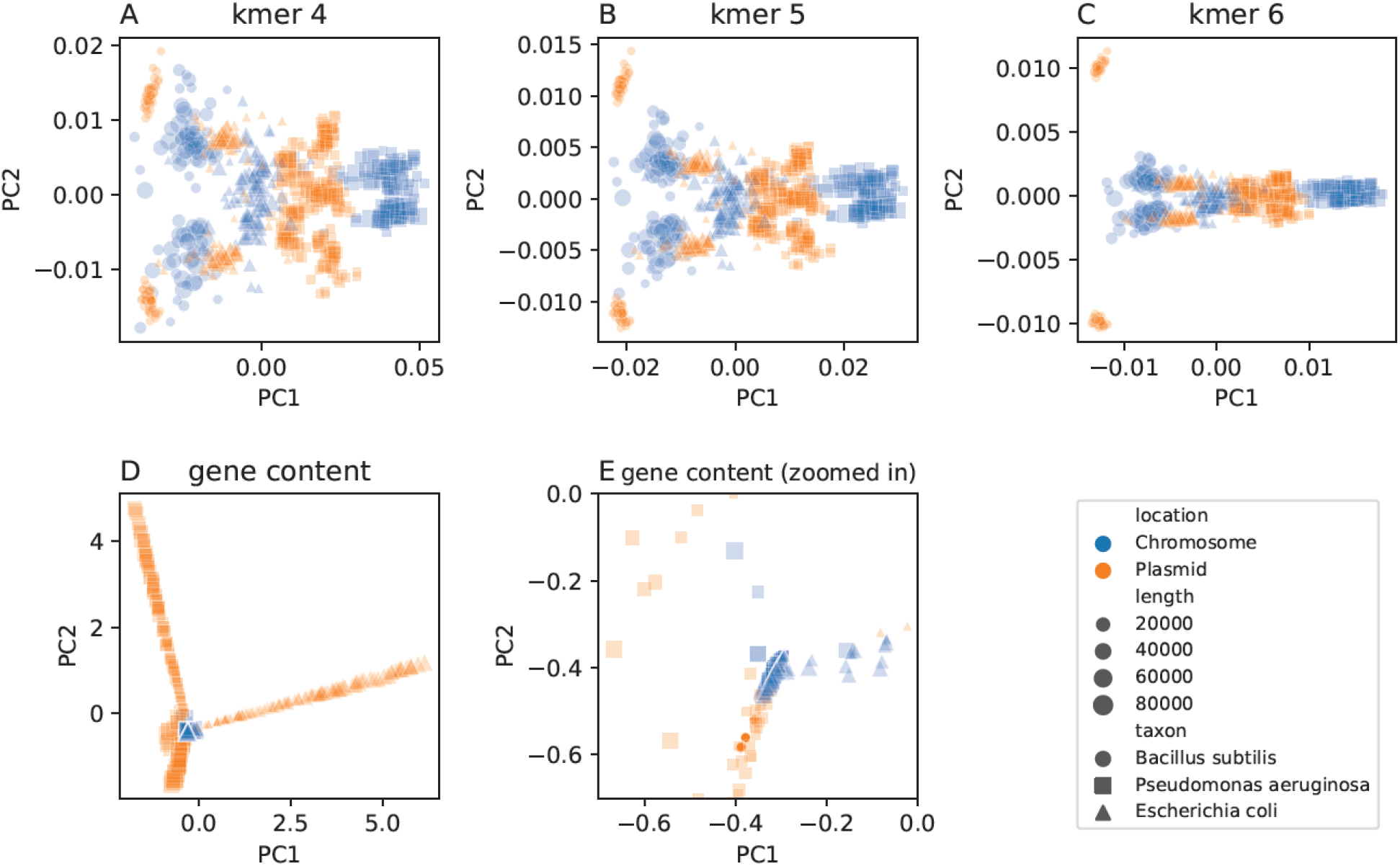
Comparisons between the discriminative power of kmer frequency profile and gene content profile in distinguishing plasmid and chromosome contigs. Three type species were used for the comparisons. For each species, 100 simulated contigs of the plasmid and chromosomal origin, respectively, were used for the PCA analysis based on the kmer 4 (**A**), kmer 5 (**B**) and kmer 6 (**C**) frequency profile and (**D**) the gene content profiles. The concentrated area of chromosomes in the figure D was zoomed in (**E**) to better visualize the extent of separation of all the chromosome contigs and the nearest plasmid contigs.

### Training dataset based on gene content profile

The downloaded 25,898 complete genomes from NCBI RefSeq database contained a total of 26,364 chromosomes (some genomes had multiple chromosomes) and 26,734 plasmids. Of these sequences, 725 chromosomes and 598 plasmids were removed from further processing due to their uncharacteristic sizes (**Supplementary Table 1**). In total, 15,000 chromosomal and 15,000 plasmid sequences were randomly selected, and then a contig with varying length from 5 Kbp to 100 Kbp (representing typical contig lengths in a genomic assembly) was randomly trimmed from each sequence (**Figure 1**). After the Prodigal gene prediction and Diamond alignment, 29,829 chromosomal and plasmid contigs were annotated using the database created in this study. The rest of the contigs had no hits and were consequently discarded. After removing contigs with < 5 unique genes, 27,093 contigs (14,641 chromosomes and 12,452 plasmids) remained for the training.

### The machine learning modeling singled out Naïve Bayes with best performance

In total, 10 models with multiple parameters were used to fit the training data (both PCA transformed and not transformed). The Naïve Bayes (NB) yielded the highest total accuracy 95.9%, and the Logistic Regression (LR) achieved the second place with an accuracy of 95.5% (**Figure 3A, Supplementary Table 2**). Because the validation dataset included almost an equal number of plasmid and chromosomal contigs, the balanced accuracies (**Figure 3B**) were nearly the same as the total accuracies. The LR had the lowest Log Loss value (0.16), and thus, it provided the highest confidence on its predicting probabilities (**Figure 3C**). In terms of sensitivity or Recall, the measure of true positive rate, the NB correctly predicted 95.1% of all the plasmid contigs (**Figure 3D**). It also had a high precision value (true positive / (true positive + false positive)) of 95.9% (**Figure 3E**), meaning that among all the predicted plasmids, 95.9% were true plasmids. Although the random forest (RF) and decision tree (DT) produced higher precision values (99.2% and 98.4%, respectively), their Recall values were very low (36.5% and 40.3%, respectively), meaning that they were too conservative in predicting a plasmid and simply ignored a big proportion of plasmids. The F score that considers both the recall and precision were calculated for all the models, and the results indicated that the NB had the best F score, 0.96 (**Figure 3F**). The indicator ROC AUC (**Figure 3G**) and ROC (**Figure 3H**) both indicated that the NB (AUC of 0.96) and LR (AUC of 0.99) were excellent, meaning that they could achieve a low false positive rate (high specificity) while maintaining a high true positive rate (high sensitivity). Considering all these indicators comprehensively, the NB had the best performance with high sensitivity and specificity, and it was chosen for the plasmid prediction tool development.

**Figure 3.**
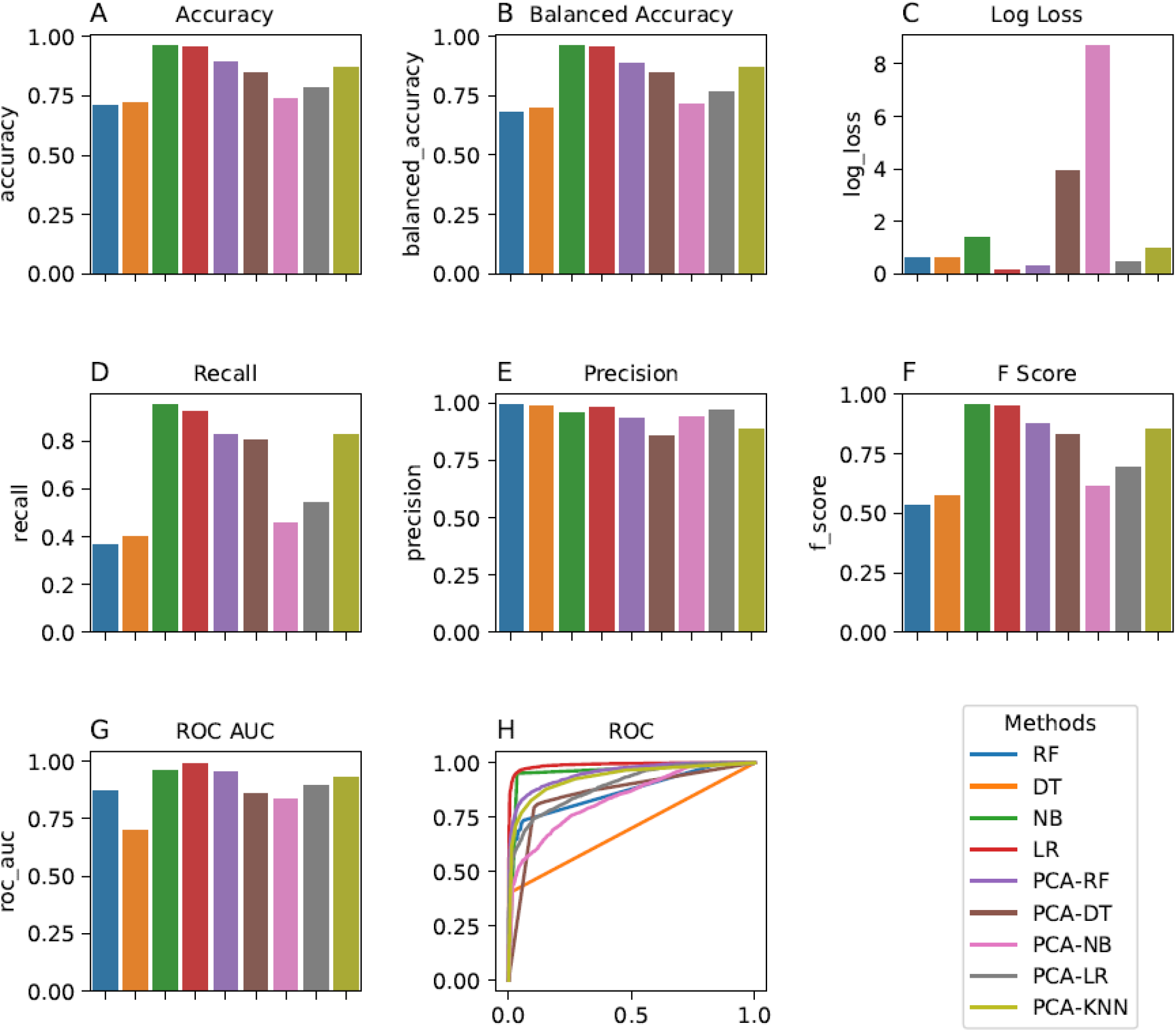
The performance evaluations of the models predicting the location (chromosome or plasmid) of contigs. The evaluations included (**A**) total accuracy, (**B**) balanced accuracy, (**C**) log loss, (**D**) recall, (**E**) precision, (**F**) F score, (**G**) AUC of ROC and (**H**) ROC. Each evaluation examined the methods of random forest (RF), decision tree (DT), naïve bayes (NB), logistic regression (LR) and K nearest neighbors (KNN). Due to the exhaustive computations required for the KNN evaluation, this evaluation could not be completed. Methods beginning with PCA- refer to the modeling using PCA transformed data.

### The PlasmidHunter outperformed other plasmid prediction tools

We generated a benchmark dataset using the genomes that were not included in the modeling (**Figure 1A**, **Supplementary Table 3**) in order to avoid introducing any biases in the benchmarking. The benchmark data included simulated contig sequences with lengths 5, 10, 20, 50 and 100 Kbps, randomly selected from the genomes. We developed a pipeline named PlasmidHunter to predict sequences based on gene content using the Naïve Bayes model (**Figure 1B**). Using the benchmark data, we compared the accuracy and speed of PlasmidHunter with those of the five recently developed tools for identify plasmids: PlasClass, PlasForest, Deeplasmid, PlasmidVerify and PlasFlow.

With the best total accuracy, PlasmidHunter outperformed all the other tools for all the selected contig length categories (5, 10, 20, 50 and 100 kb) in the five datasets with accuracies of 87.7%, 91.4%, 94.2%, 96.4% and 96.7%, respectively, (**Figure 4A, Supplementary Figure 1, Supplementary Table 4**). PlasmidHunter performed better than all the other tools on the short contigs (5 Kbp) with an accuracy of 87.7% while the accuracies of other tools ranged between 65.8% and 83.2% (**Supplementary Table 5 – 9**). Using the long contig dataset (100 Kbp), PlasmidHunter achieved an accuracy of 96.6% while the accuracies of the other tools were between 77.4% – 94.9%. Except for PlasForest, all the tools performed better with higher accuracies on longer contigs than shorter contigs. Overall, PlasmidHunter topped the list, and PlasClass, PlasFlow and PlasmidVerify were next with similar total accuracies, and they performed better than PlasForest and Deeplasmid (**Figure 4A**).

**Figure 4.**
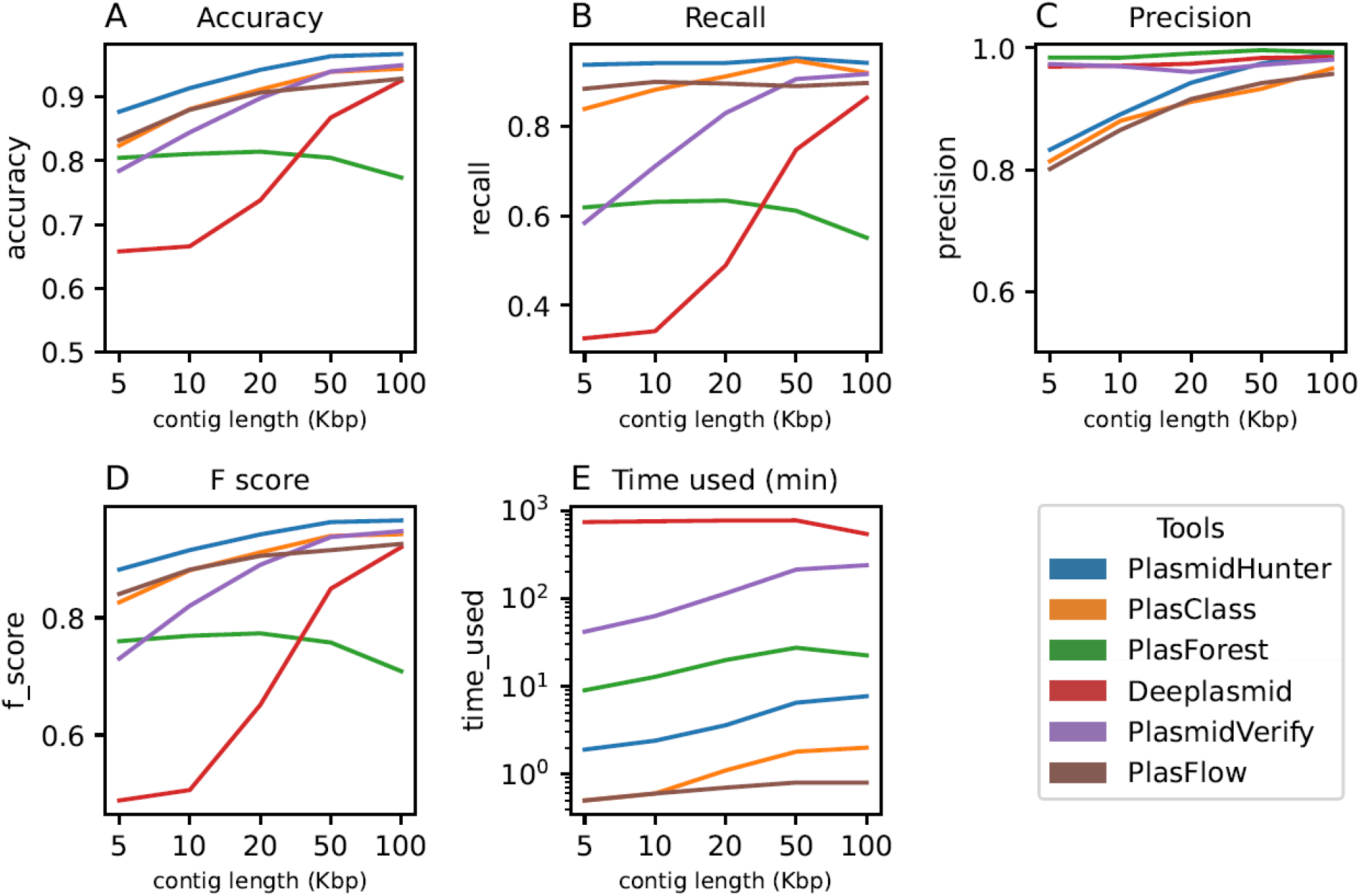
The performance evaluation of all the tools visualized on the benchmark data with different lengths. The benchmarking included evaluations on (**A**) accuracy, (**B**) recall, (**C**) precision, (**D**) F score, and (**E**) time used for running. The prediction was run on a computer with eight processors (AMD EPYC 7551, 1.2 GHz) assigned to the task, except that Deeplasmid was run on a different computer with eight processors (Intel Core i7-10510U, 1.8 GHz) because the Deeplasmid cannot be limited to use only eight processors through command. The evaluation was conducted using the package scikit-learn. The Log Loss and ROC were not shown because some tools do not output probability of prediction.

With the best recall value, PlasmidHunter again outperformed all the other tools for all contig lengths. Recall represents how sensitively the tools can detect plasmid sequences. PlasmidHunter detected 93.8%, 94.1%, 94.2%, 95.2% and 94.2% of all the plasmid sequences in the five contig length datasets (**Figure 4B**). PlasClass and PlasFlow had close recall values and were better than PlasmidVerify, PlasForest and Deeplasmid. In terms of precision, PlasForest, Deeplasmid, PlasmidVerify were the best (between 96.1% and 99.6%, **Figure 4C**), which mean that almost all the sequences predicted as plasmids were correctly predicted. However, their sensitivity or recall values were low (**Figure 4B**). Interestingly, some of the tools with the highest precision had generally a lower recall, and vice versa. In terms of F score, which is the harmonic mean of recall and precision and balances the two indicators, PlasmidHunter got the top score. PlasClass, PlasFlow and PlasmidVerify (**Figure 4D**) came next, with similar F scores that were higher than those of PlasForest and Deeplasmid.

In terms of speed, PlasmidHunter spent between 1.9 to 7.7 minutes for the five benchmark datasets. PlasFlow and PlasClass were the fastest tools, completing the task in 0.5 to 1.8 minutes. PlasmidVerify and Deeplasmid were the slowest and took much longer, up to hundreds of minutes.

## Discussion

In the past two decades, researchers in life sciences have been increasingly using HTS to investigate a wide range of important subject matters in taxonomy, evolutionary biology, ecology, agriculture, environmental and animal conservation and human health to name just a few. Consequently, the sequence databases have grown astronomically, becoming a treasure trove of information for a growing number of scientists. However, the raw sequence data that usually contains millions of reads of different lengths need to be properly checked for quality, sorted, assembled, and annotated. For example, in metagenomic data obtained through shotgun sequencing of environmental samples that usually contain many microbes, determining which reads belong to what organism is crucial for proper analyses and correct conclusions. Even whole genome sequencing of an axenic sample could produce reads that belong to different genomic compartments such as the membrane bound mitochondria, plastids and nucleus in eukaryotes or the naked plasmids and chromosomes in prokaryotes.

In addition to their chromosomes, many prokaryotes, both archaebacteria and eubacteria, carry one or more plasmids. Due to their ubiquity and mobility, plasmids are important vectors of non-essential but often beneficial traits such as virulence and AMR amongst the closely or even distantly related bacterial species. Therefore, it is critically important to develop tools that correctly determine the location/origin of the reads, plasmids or chromosomes, in the large sequence datasets. Although the latest plasmid-identifying tools have achieved some improvements, they still suffer from certain limitations. We have assessed the accuracies, recalls, precisions, F scores and speed of some of the most recently developed tools to identify plasmids, and here we introduce PlasmidHunter, a new tool that overall performs better than the other tested tools, achieving higher speeds, accuracies and recalls with contigs of various lengths.

### What are the limitations of previous plasmid-identifying tools?

Our evaluations of some of the most recent plasmid-identifying tools indicated that the use of multiple features might lead to lower accuracies. Using multiple features in ML modeling is sometimes necessary, however, multiple features introduce more assumptions, each of which might not be essential or correct. According to the heuristic Occam’s Razor in machine learning, all things being equal, simpler models can predict better than more complex models. For example, for its predictions, Deeplasmid uses both sequence signatures and gene contents—multitude of features that include GC content, homopolymer, plasmid replication origin, coding density, contig length, hit to plasmid proteins and hit to a curated Pfam database among others. Some of these features such as GC content and contig length may not be significantly and/or consistently different between chromosomes and plasmids along their entire lengths and among the short and long contigs originated from each in the assembly data with millions of reads. This might explain the lower accuracy of Deeplasmid compared to the tools with simpler/fewer features (**Supplementary Table 5**). In contrast, PlasmidHunter, PlasFlow and PlasClass all use only one type of feature, a k-mer profile or a gene content profile, and they all achieve higher accuracies and speeds than other tools that use multiple features (**Supplementary Table 4**, **Supplementary Table 7** and **Supplementary Table 9)**.

Our results also showed that the gene content profile was more discriminative than a k-mer profile for differentiating chromosomal from plasmid contigs (**Figure 2**). This may explain why PlasFlow and PlasClass, which use merely a k-mer profile as the feature, had lower accuracies than PlamidHunter.

In the benchmarking, some tools reached higher precisions for all contig lengths than others. The Deeplasmid, PlasForest and PlasmidVerify had a precision of 96.1% to 99.6% across all the benchmark datasets (**Figure 4C**). This means that almost all the contigs predicted as plasmid were true plasmid contigs. However, their low sensitivities or recall values, especially for short contigs (down to 36.2%, **Figure 4B**), mean that they detect a small percentage of all the plasmid contigs in the dataset. As a result, their F scores were also much lower than the others (**Figure 4D**). In comparison, PlasmidHunter had a more balanced recall and precision than all the other tools.

### What are PlasmidHunter’s strengths and weaknesses?

Compared to the previous top plasmid-identifying tools and based on the benchmarking using datasets of sequences with different lengths, PlasmidHunter achieved the best total accuracies and recalls or sensitivities for all contig lengths (**Figure 4A, Figure 4B**). On the downside, PlasmidHunter showed a lower precision than some of the other tools, especially for short contigs (83.3% and 89.1% for 5 kbp and 10 Kbp contigs, respectively), but due to its higher recall values for all contig lengths, it achieved a better balance (**Figure 4B, Figure 4C**). F score, the harmonic mean of the recall and the precision, serves as a measure of balance between the recall and precision. PlasmidHunter had the highest F scores for all lengths (**Figure 4D**). In terms of speed, PlasmidHunter completed running the test data within 1.9 to 7.7 minutes and ranked the third among all the tools (**Figure 4E**).

## Conclusions

Here, we have made rigorous comparisons between recently-published plasmid-prediction tools using independent benchmark datasets of contigs with different lengths. We showed that the use of multiple and complex features does not necessarily result in higher accuracy in modeling. Our study also provides useful insights into feature selection as well as ML algorithms for modeling in sequence classifications. Finally, we present our tool, PlasmidHunter, achieving the highest accuracy, recall and F score.

## Materials and Methods

### Database preparation

The NCBI prokaryotic genome databases were searched for with the filtration assembly level of ‘complete’ (https://www.ncbi.nlm.nih.gov/genome/browse#!/prokaryotes/, and the results were downloaded as metadata. As of October 26, 2021 there were 25,898 complete genomes of prokaryotes. A custom Python script was used to download all the protein sequences of the genomes using the URLs in the metadata with multiple processes. The protein sequences were then clustered using MMseqs2 (version 13.45111) [22] with sequence identity cutoff of 0.4, alignment coverage of 0.8 and --cov-mode of 0 (bidirectional). The clusters with < 5 sequences were removed. The representative sequences of the remaining 0.96 million clusters (i.e., one representative sequence from each cluster) were used as a database for gene content profiling.

### Gene content annotation

Gene content profiling was conducted by alignment to the sequences in the protein database. First, genes were predicted using Prodigal (version 2.6.3) [23] in the meta or anonymous mode and allowing genes to run off edges. The protein sequences were then used to align against the database using BLASTp mode of DIAMOND (version 2.0.15.153) [24]. Query coverage of 80%, protein sequence identity of 40% and E value of 1e-5 were used to filter the hits (-max-target-seqs 1 --max-hsps 1 --evalue 1e-5 --id 40 --query-cover 80). The protein IDs of the hits with the highest alignment scores were assigned to the queries. The gene content profile was then summarized into a data frame in Python.

### Comparing the discriminatory power of gene content-based and k-mer-based methods

Three bacterial type strains, *Escherichia coli* (GCA_000008865.2), *Bacillus subtilis* (GCA_009363835.1) and *Pseudomonas aeruginosa* (GCA_019738995.1) were used for the comparison. Genome sequences were downloaded using ncbi-genome-download (version 0.3.0) and 100 simulated contigs of chromosomes and plasmids, respectively, with random lengths between 5 and 100 Kbp were generated for each genome. The gene content profile of the simulated contigs was generated as described above. The k-mer frequency profile (k = 4, 5 and 6, respectively) of the simulated contigs was generated using custom codes with

Python package collections.Counter and Pandas. The gene content and k-mer frequency profiles were then compared by dimensionality reduction with principal component analysis (PCA). The Python package sklearn.decomposition.PCA was used to conduct the PCA analysis with two components. The outputs were then visualized with a Python package seaborn (version 0.11.2).

### Training data set for modeling

The gff files of the 25,898 complete prokaryotic genomes from NCBI RefSeq database were downloaded using the URLs in the metadata and parsed using the Python package BCBio in a multiprocessing mode. For each gff record, the sequence length, location (plasmid or chromosome) and a list of protein IDs in the original order were extracted and saved in a json file. To exclude any false annotation of sequence location, the sequence was defined as chromosomal if the annotation was ‘chromosome’ and the size was > 900 Kbp, and as plasmid if the annotation was ‘plasmid’ and the size was < 600 Kbp.

For the training dataset, 15,000 plasmid and 15,000 chromosome sequences were randomly selected and the rest of the sequences were set aside for model testing and evaluation. The DNA sequences of genomes were downloaded using the URLs in the metadata. For each selected sequence, a fragment with a random start point and a random length (from 5 Kbp to 100 Kbp, representing typical contig lengths in a genomic assembly) was selected. If the sequence length was less than the random length, the whole sequence would be used. The simulated contigs would then be annotated as mentioned above and a feature table of gene presence and absence of all the simulated contigs would be generated in Python Pandas, using the representative protein IDs as features.

### Machine learning modeling

The Python package scikit-learn (version 1.0.2) was used for the modeling. The training data were first split into training dataset (75%) and validation dataset (25%) using the function train_test_split. For the modeling, the following algorithms were tested to fit the models: Decision Tree (with maximum depth of 5, 10, 15 and 20); Random Forest (with maximum depth of 5, 10, 15 and 20); Naïve Bayes; Logistic Regression; Support Vector Machine (with regularization 0.1, 1 and 10); and K Nearest Neighbors (n=7). The validation dataset was, then, used to calculate the accuracy of the classifications. Meanwhile, the training data were transformed with PCA transformation (with PC number 30), and the transformed data were subjected to the same processes as mentioned above.

### Model validation

The validation dataset (25% of training data) was held back to be used for the model validation. For each model, the prediction scores, hence the probabilities of each class, were calculated using the function predict_proba. A custom function accepting inputs of true classification (y_true) and prediction scores (y_score) was used to evaluate each model’s performance with the package sklearn.metrics. The total accuracy was calculated with the function accuracy_score. The Log Loss was calculated with the function log_loss. The recall (true positive rate) and precision (true positive / (true positive + false positive)) were calculated with the function recall_score and precision_score, respectively. An F score was based on the harmonic mean of the recall and precision. Finally, an area under curve (AUC) of the receiver operating characteristic (ROC) curve was calculated with the function roc_auc. The best model was ‘dumped’ as a local binary file using pickle.

### Benchmark data for model test and evaluation

As mentioned above, 15,000 chromosome and plasmid sequences, respectively, from the 25,898 complete prokaryotic genomes from RefSeq database, were used for the modeling and validation. The remaining data was held back from the modeling for testing and unbiased evaluation. The balanced datasets including equal number of chromosome and plasmid sequences (3,000 each) were randomly selected. Contigs of different lengths, including 5, 10, 20, 50 and 100 Kbps, were randomly simulated from each sequence and were used for the gene content profiling and prediction. The contigs were annotated using DIAMOND search against the database described above. The gene content profile was then processed to ensure that it had the same features that were used/included in the modeling data. The model was loaded using pickle and was used to predict the feature data. The performance of the model on the data of different sequence lengths was then evaluated with the package sklearn.metrics as mentioned in the model validation.

### Construction of the gene content-based plasmid prediction pipeline

A pipeline named PlasmidHunter was developed using the validated model to predict plasmid sequences from input contigs. First, the input contigs are filtered to remove short sequences (< 5 Kbp). Prodigal (version 2.6.3) is used to predict gene and protein sequences. Diamond (version 2.0.15.153) is used to search the protein sequences against the custom database. A gene content profile is generated using the alignment results based on the features used in the modeling step. The gene content profile is then used as features for prediction using the Python package sklearn (version 1.0.2) and the naïve bayes model.

### Benchmarking of multiple plasmid predictors

Six tools, including PlasmidHunter, PlasClass, PlasFlow, PlasmidVerify, PlasForest and Deeplasmid were tested using the benchmark datasets with different contig lengths. All the tools were run following the manuals provided, on a high performance computer (HPC) with eight processors (AMD EPYC 7551, 1.2 GHz) assigned to the tasks, except that Deeplasmid was run on a different computer with eight processors (Intel Core i7-10510U, 1.8 GHz) because the Deeplasmid cannot be limited to use only eight processors through command line. The running was timed using the Python package, time. The outputs were parsed to retrieve the prediction of each contig and the corresponding probability if any. A Python package sklearn.metrics was used to evaluate the prediction results in terms of total accuracy, balanced accuracy, recall, precision, F score and AUC ROC. A Python package seaborn was used to visualize the evaluation results.

### Availability of the scripts and the benchmark data sets of this study

All the scripts for data processing and analysis and the benchmark datasets with different contig lengths have been deposited on GitHub (https://github.com/tianrenmaogithub/PlasmidHunter).

### Availability of PlasmidHunter source code and web server

The source code of PlasmidHunter is available on GitHub for plasmid prediction analysis (https://github.com/tianrenmaogithub/PlasmidHunter). We have also built a web server (https://colab.research.google.com/github/tianrenmaogithub/PlasmidHunter/blob/main/PlasmidHunter.ipynb) for users to run the analysis online.

## Ethics approval and consent to participate

Not applicable

## Consent for publication

Not applicable

## Availability of data and materials

The benchmark datasets with different contig lengths have been deposited on GitHub (https://github.com/tianrenmaogithub/PlasmidHunter).

## Author contributions

B.I. and R.T. conceptualized and designed the project; R.T. conducted the data analysis, code writing and pipeline development. R.T. and B.I. wrote the manuscript. All authors have read and agreed to the published version of the manuscript.

## Acknowledgments

The authors acknowledge the Food and Drug Administration (FDA) of the U.S. Department of Health and Human Services (HHS) for the support.

