## supplementary information for "PlasmidHunter: Accurate and fast prediction of plasmid sequences using gene content profile and machine learning"

**Supplementary Table 1.** Summary of the confirmed chromosomes and plasmids for the modeling based on the annotation from NCBI database and the sizes. The sequences determined as chromosome or plasmid were included in the subsequent analaysis.

| **Annotation** | **Size** | **Determined as** | **Count** |
|---|---|---|---|
| chromosome | > 900 Kbp | chromosome | 25651 |
| chromosome | <= 900 Kbp | not sure | 713 |
| plasmid | < 600 Kbp | plasmid | 26136 |
| plasmid | >= 600 Kbp | not sure | 598 |
| extrachrom | | not sure | 7 |
| genomic | | not sure | 5 |

**Supplementary Table 2.** The evaluation of the performance of models predicting the location (chromosome or plasmid) of contigs. The evaluation was based on the model predictions on validation data using the package scikit-learn. RF = random forest; DT = decision tree; NB = naïve bayes; LR = logistic regression. Methods beginning with principal component analysis (PCA) means the modeling using PCA transformed data. Recall was calculated as TP/(TP+FN) and Precision was calculated as TP/(TP+FP), where TP is true positive, FN is false negative, and FP is false positive. F1 Score was calculated as 2*(Recall * Precision) / (Recall + Precision).

| **Methods** | **Accuracy** | **Balanced accuracy** | **Log loss** | **Recall** | **Precision** | **F score** | **ROC AUC** |
|---|---|---|---|---|---|---|---|
| RF | 70.7% | 68.1% | 0.60 | 36.5% | 99.2% | 0.53 | 0.87 |
| DT | 72.2% | 69.9% | 0.60 | 40.3% | 98.4% | 0.57 | 0.70 |
| ***NB*** | ***95.9%*** | ***95.9%*** | ***1.41*** | ***95.1%*** | ***95.9%*** | ***0.96*** | ***0.96*** |
| LR | 95.5% | 95.3% | 0.16 | 92.2% | 97.9% | 0.95 | 0.99 |
| PCA-RF | 89.1% | 88.6% | 0.29 | 82.4% | 93.1% | 0.87 | 0.95 |
| PCA-DT | 84.9% | 84.6% | 3.93 | 80.4% | 85.9% | 0.83 | 0.86 |
| PCA-NB | 73.7% | 71.6% | 8.68 | 45.7% | 94.1% | 0.62 | 0.84 |
| PCA-LR | 78.2% | 76.4% | 0.45 | 54.3% | 96.8% | 0.70 | 0.90 |
| PCA-KNN | 87.1% | 86.8% | 0.99 | 82.6% | 88.6% | 0.85 | 0.93 |

**Supplementary Table 3.** The statistics of the benchmark data. The contigs of specified lengths were randomly simulated from 3,000 plasmids and 3,000 chromosomes that were not included in the modeling. Each data set has equal number of plasmid and chromosome sequences.

| **Bechmark data ID** | **Contig length (Kbp)** | **Number of sequences *** | **Total length (Mbp)** |
|---|---|---|---|
| 1 | 5 | 6000 | 30 |
| 2 | 10 | 5230 | 52.3 |
| 3 | 20 | 4746 | 94.92 |
| 4 | 50 | 3520 | 176 |
| 5 | 100 | 1996 | 199.6 |

* The total number of sequences were less than 6000 in some data sets because some plasmids were shorter than the required lengths and were thus not included.

**Supplementary Table 4.** The evaluation of PlasmidHunter's performance on the benchmark data with different lengths. The prediction was run on a computer with eight processors (AMD EPYC 7551, 1.2 GHz) assigned to the task. The evaluation was conducted using the package scikit-learn. Recall was calculated as TP/(TP+FN) and Precision was calculated as TP/(TP+FP), where TP is true positive, FN is false negative, and FP is false positive. F1 Score was calculated as 2*(Recall * Precision) / (Recall + Precision).

| Contig length (Kbp) | Accuracy | Balanced accuracy | Log loss | Recall | Precision | F score | ROC AUC | Time used (min) |
|---|---|---|---|---|---|---|---|---|
| 5 | 87.70% | 87.80% | 4.257 | 93.80% | 83.30% | 88.20% | 87.80% | 1.9 |
| 10 | 91.40% | 91.40% | 2.983 | 94.10% | 89.10% | 91.50% | 91.40% | 2.4 |
| 20 | 94.20% | 94.20% | 1.993 | 94.20% | 94.30% | 94.20% | 94.20% | 3.6 |
| 50 | 96.40% | 96.40% | 1.257 | 95.20% | 97.40% | 96.30% | 96.40% | 6.5 |
| 100 | 96.70% | 96.70% | 1.142 | 94.20% | 99.20% | 96.60% | 96.70% | 7.7 |

**Supplementary Table 5.** The evaluation of Deeplasmid (version of Feb. 10, 2022) performance on the benchmark data with different lengths. The prediction was run on a different computer with eight processors (Intel Core i7-10510U, 1.8 GHz) because the Deeplasmid cannot be limited to use only eight processors in the other computer with 128 processors. Note that CPU mode rather than GPU mode of Deeplasmid was used in this running. The evaluation was conducted using the package scikit-learn. Recall was calculated as TP/(TP+FN) and Precision was calculated as TP/(TP+FP), where TP is true positive, FN is false negative, and FP is false positive. F1 Score was calculated as 2*(Recall * Precision) / (Recall + Precision).

| Contig length (Kbp) | Accuracy | Balanced accuracy | Log loss | Recall | Precision | F score | ROC AUC | Time used (min) |
|---|---|---|---|---|---|---|---|---|
| 5 | 65.8% | 65.8% | 1.340 | 32.6% | 96.9% | 48.8% | 86.2% | 742 |
| 10 | 66.6% | 66.6% | 1.901 | 34.2% | 97.1% | 50.6% | 88.8% | 758 |
| 20 | 73.8% | 73.8% | 1.971 | 48.9% | 97.4% | 65.1% | 92.3% | 773 |
| 50 | 86.8% | 86.8% | 1.278 | 74.8% | 98.4% | 85.0% | 95.6% | 776 |
| 100 | 92.5% | 92.5% | 0.623 | 86.4% | 98.5% | 92.0% | 97.8% | 543 |

**Supplementary Table 6.** The evaluation of PlasmidVerify (version of April 30, 2020) performance on the benchmark data with different lengths. The prediction was run on a computer with eight processors (AMD EPYC 7551, 1.2 GHz) assigned to the task. The evaluation was conducted using the package scikit-learn. The log loss and ROC AUC were not calculated because PlasmidVerify outputs likelihood ratios rather than probabilities of predictions. Recall was calculated as TP/(TP+FN) and Precision was calculated as TP/(TP+FP), where TP is true positive, FN is false negative, and FP is false positive. F1 Score was calculated as 2*(Recall * Precision) / (Recall + Precision).

| Contig length (Kbp) | Accuracy | Balanced accuracy | Recall | Precision | F score | Time used (min) |
|---|---|---|---|---|---|---|
| 5 | 78.42% | 78.42% | 58.43% | 97.33% | 73.03% | 41.8 |
| 10 | 84.44% | 84.44% | 71.09% | 96.97% | 82.04% | 63.2 |
| 20 | 89.78% | 89.78% | 82.98% | 96.05% | 89.03% | 114.3 |
| 50 | 93.98% | 93.98% | 90.57% | 97.20% | 93.76% | 213.6 |
| 100 | 94.94% | 94.94% | 91.68% | 98.07% | 94.77% | 239.5 |

**Supplementary Table 7.** The evaluation of PlasFlow (version 1.1.0, August 15, 2018) performance on the benchmark data with different lengths. The prediction was run on a computer with one processors (AMD EPYC 7551, 1.2 GHz) assigned to the task, because PlasmidFlow does not have multiple processing mode. Threshold of 0.5 was used for classification to eliminate unclassified prediction. The evaluation was conducted using the package scikit-learn. The log loss and ROC AUC were not calculated because PlasFlow does not output probability of predictions plasmid or chromosome. Recall was calculated as TP/(TP+FN) and Precision was calculated as TP/(TP+FP), where TP is true positive, FN is false negative, and FP is false positive. F1 Score was calculated as 2*(Recall * Precision) / (Recall + Precision).

| Contig length (Kbp) | Accuracy | Balanced accuracy | Recall | Precision | F score | Time used (min) |
|---|---|---|---|---|---|---|
| 5 | 83.23% | 83.23% | 88.40% | 80.12% | 84.06% | 0.5 |
| 10 | 87.97% | 87.97% | 89.94% | 86.53% | 88.21% | 0.6 |
| 20 | 90.67% | 90.67% | 89.55% | 91.59% | 90.56% | 0.7 |
| 50 | 91.76% | 91.76% | 88.98% | 94.22% | 91.53% | 0.8 |
| 100 | 92.84% | 92.84% | 89.68% | 95.72% | 92.60% | 0.8 |

**Supplementary Table 8.** The evaluation of PlasForest (version of October 14, 2021) performance on the benchmark data with different lengths. The prediction was run on a computer with eight processors (AMD EPYC 7551, 1.2 GHz) assigned to the task. The evaluation was conducted using the package scikit-learn. The log loss and ROC AUC were not calculated because PlasForest does not output probability of prediction. Recall was calculated as TP/(TP+FN) and Precision was calculated as TP/(TP+FP), where TP is true positive, FN is false negative, and FP is false positive. F1 Score was calculated as 2*(Recall * Precision) / (Recall + Precision).

| Contig length (Kbp) | Accuracy | Balanced accuracy | Recall | Precision | F score | Time used (min) |
|---|---|---|---|---|---|---|
| 5 | 80.45% | 80.45% | 61.90% | 98.41% | 76.00% | 9 |
| 10 | 81.05% | 81.05% | 63.14% | 98.39% | 76.92% | 12.8 |
| 20 | 81.42% | 81.42% | 63.42% | 99.08% | 77.34% | 19.9 |
| 50 | 80.45% | 80.45% | 61.14% | 99.63% | 75.77% | 27.5 |
| 100 | 77.35% | 77.35% | 55.11% | 99.28% | 70.88% | 22.5 |

**Supplementary Table 9.** The evaluation of PlasClass (version 0.1.1, November 1, 2021) performance on the benchmark data with different lengths. The prediction was run on a computer with eight processors (AMD EPYC 7551, 1.2 GHz) assigned to the task. The evaluation was conducted using the package scikit-learn. Recall was calculated as TP/(TP+FN) and Precision was calculated as TP/(TP+FP), where TP is true positive, FN is false negative, and FP is false positive. F1 Score was calculated as 2*(Recall * Precision) / (Recall + Precision).

| Contig length (Kbp) | Accuracy | Balanced accuracy | Log loss | Recall | Precision | F score | ROC AUC | Time used (min) |
|---|---|---|---|---|---|---|---|---|
| 5 | 82.37% | 82.37% | 0.42 | 83.90% | 81.40% | 82.63% | 90.44% | 0.5 |
| 10 | 88.09% | 88.09% | 0.30 | 88.18% | 88.02% | 88.10% | 94.71% | 0.6 |
| 20 | 91.15% | 91.15% | 0.23 | 91.15% | 91.15% | 91.15% | 96.76% | 1.1 |
| 50 | 93.92% | 93.92% | 0.18 | 94.66% | 93.28% | 93.97% | 98.13% | 1.8 |
| 100 | 94.39% | 94.39% | 0.68 | 91.98% | 96.63% | 94.25% | 97.94% | 2 |

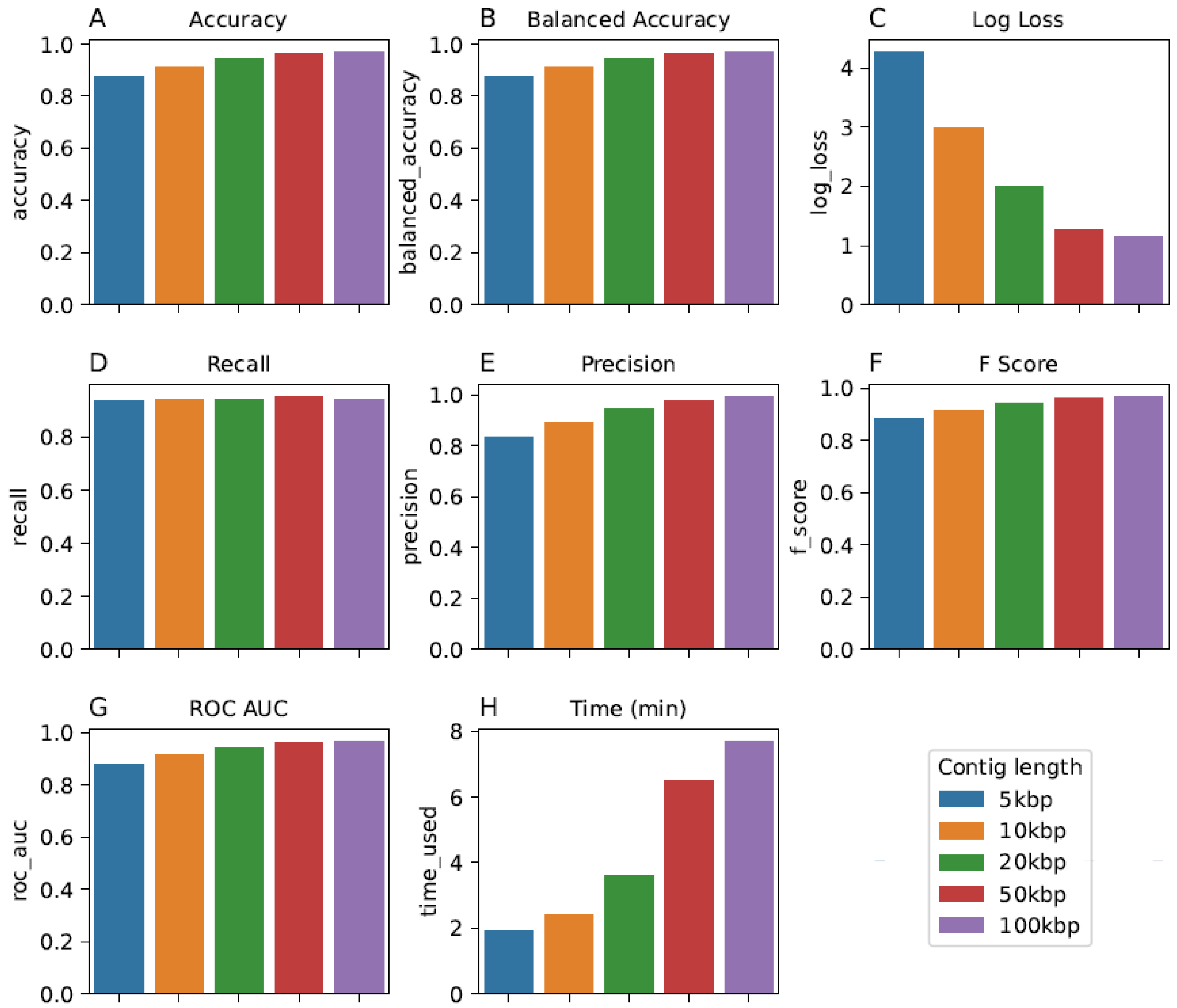

**Supplementary Figure 1.** The visualized evaluation of PlasmidHunter's performance on the benchmark data with different lengths. The prediction was run on a computer with eight processors (AMD EPYC 7551, 1.2 GHz) assigned to the task. The evaluation was conducted using the package scikit-learn.
